# Comprehensive profiling of the fission yeast transcription start site activity during stress and media response

**DOI:** 10.1101/281642

**Authors:** Malte Thodberg, Axel Thieffry, Jette Bornholdt, Mette Boyd, Christian Holmberg, Ajuna Azad, Christopher T. Workman, Yun Chen, Karl Ekwall, Olaf Nielsen, Albin Sandelin

**Affiliations:** The Bioinformatics Centre, Department of Biology and Biotech Research and Innovation Centre, University of Copenhagen, Denmark; Cell cycle and genome stability group, Department of Biology, University of Copenhagen, Denmark; Department of Biotechnology and Biomedicine, Technical University of Denmark, Denmark; Department of Biosciences and Nutrition, Karolinska Institute, Sweden

## Abstract

Fission yeast, *Schizosaccharomyces pombe,* is an attractive model organism for transcriptional and chromatin biology research. Such research is contingent on accurate annotation of transcription start sites (TSSs). However, comprehensive genome-wide maps of TSSs and their usage across commonly applied laboratory conditions and treatments for *S. pombe* are lacking. To this end, we profiled TSS activity genome-wide in *S. pombe* cultures exposed to heat shock, nitrogen starvation, hydrogen peroxide and two commonly applied media, YES and EMM2, using Cap Analysis of Gene Expression (CAGE). CAGE-based annotation of TSSs is substantially more accurate than existing PomBase annotation; on average, CAGE TSSs fall 50-75 bp downstream of PomBase TSSs and co-localize with nucleosome boundaries. In contrast to higher eukaryotes, *S. pombe* does not show sharp and dispersed TSS distributions. Our data recapitulate known *S. pombe* stress expression response patterns and identify stress- and media-responsive alternative TSSs. Notably, alteration of growth medium induces changes of similar magnitude as some stressors. We show a link between nucleosome occupancy and genetic variation, and that the proximal promoter region is genetically diverse between *S. pombe* strains. Our detailed TSS map constitute a central resource for *S. pombe* gene regulation research.

## Introduction

Yeast cells have been central models for understanding eukaryotic gene regulation. Historically, baker’s yeast (*Saccharomyces cerevisiae*) has been the unicellular model of choice, but the remotely related fission yeast *Schizosaccharomyces pombe* has in many ways turned out to be a more relevant model for mammalian cells (1). The architecture of *S. pombe* and human chromosomes are similar, featuring large repetitive centromeres and regions of RNAi-dependent heterochromatin (2). *S. pombe* also utilizes histone modifications and chromatin remodelling enzymes similar to multicellular eukaryotes (2). On the other hand, both yeast types have highly related signal transduction pathways responding to environmental stress (3).

A central challenge for cells is to react to changing environments through the regulated activation of gene transcription. The most commonly studied gene regulation responses in yeast cells are to environmental stress through chemicals (e.g. hydrogen peroxide, sorbitol, cadmium), physical conditions (e.g. heat), or changes in available nutritional substances (nitrogen starvation, change of growth media, etc.). Previous work on stress response in *S. pombe* and *S. cerevisiae* has focused on the distinction between a general environmental stress response (Core Environmental Stress Response, CESR) versus specific response to individual types of stress (Specific Environmental Stress Response, SESR). CESR is comprised of metabolic genes related to carbohydrate metabolism and genes involved in protein stability such as anti-oxidants, proteases and heat shock proteins, while SESR is comprised of genes with more specific functions related to the given type of stress (4–7).

In *S. pombe,* the timing and maintenance of the CESR is controlled by a conserved signal transduction pathway that ultimately activates the Sty1 protein kinase (*sty1*) (8), a member of the stress-activated MAP kinase family (SAPK family), which also includes HOG1 from *S. cerevisiae* and human *MAPK14* and cJun-N terminal kinase *(MAPK8).* Sty1 in turn activates key transcription factors such as Atf1, Pap1 and Hsf1 (8–10).

Stress-specific gene regulation has been studied by comparing genome-wide changes in expression profiles in wild-type cells and mutants lacking key signaling components or transcription factors (4, 11). While some types of stress (for example heat shock) induce a quick and transient response, hydrogen peroxide and alkylating agents induce a more lengthy response (4). Furthermore, both stress exposure and environmental cues, such as nutritional composition, modulate the growth rate and size at which fission yeast cells enter mitosis (12). This size regulation occurs partly through the Sty1 SAPK and also via the TOR (target of rapamycin) pathway (13, 14). Extreme nutritional stresses, in particular nitrogen starvation, induces cells to enter sexual development, a process which is intertwined with the CESR (15).

Accurate maps of TSSs and their activity have been instrumental in understanding gene regulation and core promoter activity, as well as the evolution of gene regulation in mammals, birds, insects and plants (e.g. (16–21)). Such data sets have also been highly beneficial in deciphering the relationship between chromatin and transcription initiation (e.g. (22–25)). Due to its importance as a model organism in chromatin biology and stress response research, it is surprising that no comprehensive maps of transcription start sites (TSSs) across different cellular states have been reported for *S. pombe.*

Cap Analysis of Gene Expression (CAGE), based on sequencing the first 20-30 bp of capped, full-length cDNAs (26) is arguably the most used technique for locating TSSs and their transcriptional activity genome-wide (27). Such so-called CAGE tags can, when mapped to a reference genome, identify TSSs with single nucleotide resolution and quantify their level of transcription, as the number of CAGE tags mapping to a location is proportional to the concentration of the originating mRNA (Fig. 1A). CAGE tags can thus be used for expression profiling, but on TSS rather than gene level. In this sense, it is complementary to RNA-Seq, which has the advantage that splicing can be assessed but, on the other hand, is not precise in locating TSSs. Previous studies have shown that RNA-Seq and CAGE have comparable accuracy in terms of expression profiling (28). Because CAGE is not contingent on existing gene models, it can both refine existing gene models and locate novel TSSs, both within and outside of known genes. In eukaryotes, alternative TSS usage is common (29): for instance, in human and mouse >50% of genes have two or more distinct TSSs, many of which are used in a tissue-or context-specific manner (21). Such alternative TSSs are interesting as they may confer additional, independent regulatory inputs for genes and/or change the protein-coding potential of the resulting mRNA, for instance by excluding exons coding for protein domains (30, 31).

**Fig 1.**
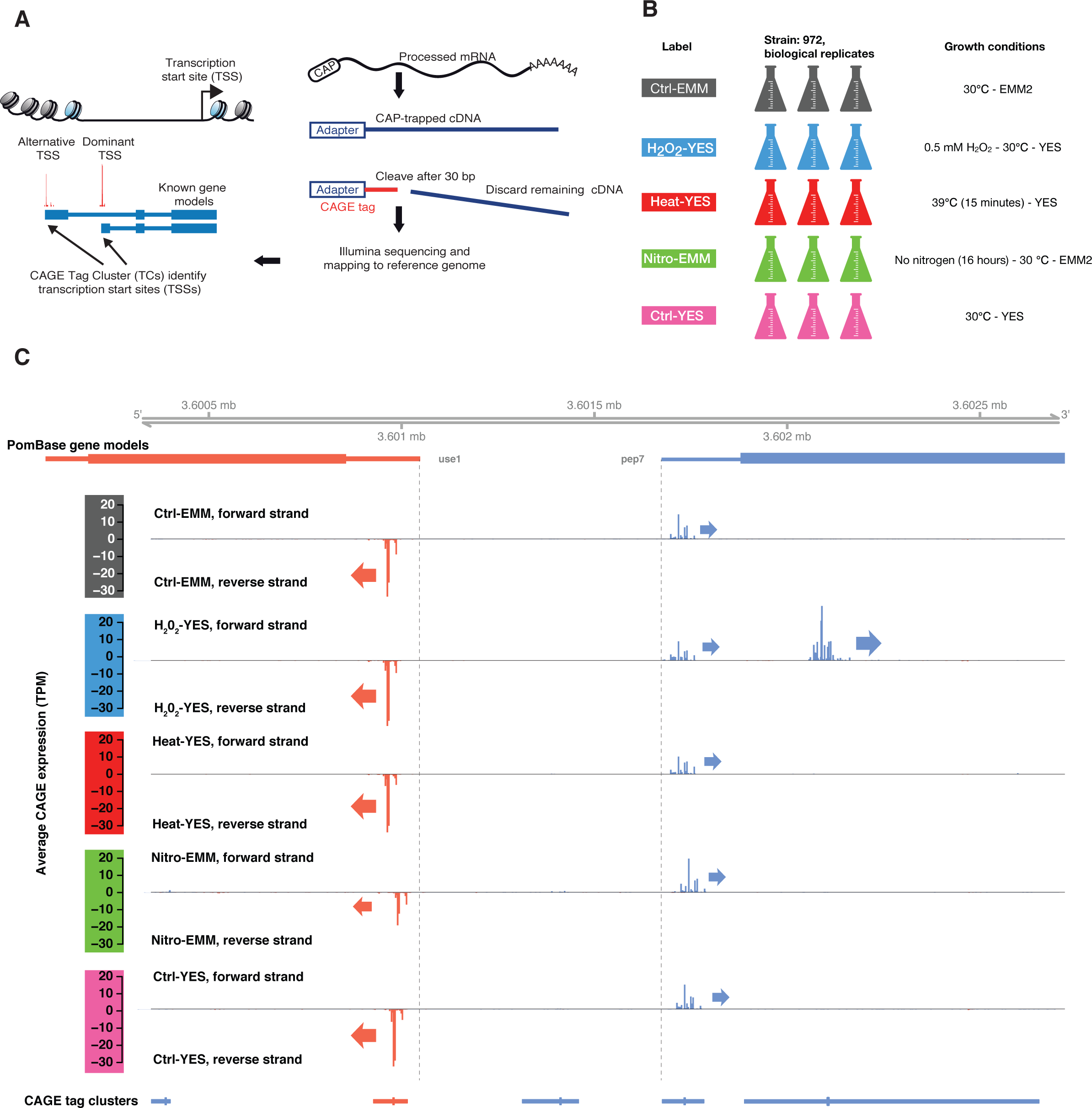
Overview of CAGE experiment. **A: Schematic overview of the CAGE protocol.**Capped full-length cDNAs are isolated from total RNA through cap-trapping, followed by cleaving off the first 30 bp, which are sequenced and mapped back to the reference genome, identifying TSS locations and their relative usage. Such 30 bp reads are referred to as CAGE tags. Nearby CAGE tags on the same strand are grouped into tag clusters.
**B: Experimental design.** Five sets of biological *S. pombe* triplicates were prepared, covering three stressors (heat shock, hydrogen peroxide stress and nitrogen starvation) and two common growth media: Edinburgh Minimal Media (EMM2) and Yeast Extract (YES). Labels for samples/libraries are shown in the left column. Growth conditions are shown in the right column.
**C: Genome-browser example of CAGE defined TSS landscape of the *pep7* gene locus.** X-axis represents the genome. Panels represent different types of data mapped to the genome. Data on forward strand is indicated by blue color, reverse strand by red. The first and second tracks show PomBase gene models; dotted vertical lines show PomBase-defined TSSs. The third to seventh tracks show the average pooled CAGE signal from each experimental group (indicated by color code to the left) as bar plots where the Y-axis shows CAGE expression (TPM-normalized number of 5’-ends of CAGE tags at a given bp), where positive values and blue color indicate forward strand CAGE-tags, and negative values and red color indicate reverse strand CAGE-tags. Arrows show the direction of transcription for highly used TSSs. The last track shows the location of clusters of nearby CAGE tags, where thicker lines indicate the TSS peaks, i.e. the position with the highest CAGE signal. We identified ubiquitously expressed TSS for *use1* and *pep7* located just downstream of the PomBase gene model-defined TSSs, but also a novel *pep7* alternative TSS only active in H_2_O_2_-YES. Also, see **Fig. S1B-E** for additional genome browser examples.

Since stress response is highly studied in *S. pombe,* having accurate and genome-wide TSS maps for stress states would be beneficial for understanding their gene regulation and associated processes. Here, we used CAGE to define a TSS atlas of unprecedented resolution and scope for *S. pombe,* across three environmental stressors (heat shock, nitrogen starvation and hydrogen peroxide stress) and two commonly used growth media, Edinburgh Minimal Medium (EMM2) and Yeast Extract Medium with supplements (YES). We show that this atlas substantially expands and refines current state-of-the-art *S. pombe* TSS annotation, allowing for more detailed interpretation of a range of processes, including nucleosome positioning, histone modification and transcription levels. Additionally, our CAGE-based annotation allows the analysis of stress-specific and shared stress response regulation and growth media adaptation, and enables refined analysis of *S. pombe* genetic variation data.

## Materials and Methods

### Culture conditions and RNA preparation

Triplicate cultures of the wild-type strain *h^-S^* (972) were grown at 30°C in either EMM2 (Edinburgh minimal medium) or YES (yeast extract with supplements) to a density of 5 X 10^6^ cells/ml (32). For nitrogen starvation, cells were transferred by vacuum filtration from EMM2 to EMM2 without ammonium chloride and incubated for 16 hours at 30°C. Heat shock was imposed by transferring YES cultures to 39°C for 15 minutes. Oxidative stress was inflicted by treating YES cultures with 0.5 mM H_2_O_2_ for 15 minutes at 30°C. Total RNA was extracted from 10^8^ cells. In brief, cells were harvested by 5 minutes of centrifugation at 3000 rpm. Pellets were resuspended in TES (10 mM TrisHCl pH7.5, 10 mM EDTA and 0.5% SDS) and transferred to 65°C preheated acidic phenol-chloroform (Sigma P-1944). After 1 hour of incubation at 65°C with mixing every 10 minutes, RNA was extracted with chloroform-isoamyl alcohol (Sigma C-0549), ethanol precipitated and re-suspended in water. All RIN-values were above 8.8. As CAGE requires a certain amount of input RNA concentration, RNA from nitrogen-starved cultures was additionally concentrated by vacuum centrifuge (reported RIN-scores are after concentration). See **Table S1** for an overview of libraries.

### CAGE analysis and mapping

CAGE libraries were prepared from the 15 yeast cultures as in (26), using 5 μg total RNA as starting material. Libraries were run individually with the following four barcode: #2(CTT), #3 (GAT), #4 (CACG) and #8 (ATC). Prior to sequencing, four CAGE libraries with different barcodes were pooled and applied to the same sequencing lane. Sequencing of the libraries was performed on a HiSeq2000 instrument from Illumina at the National High-Throughput DNA Sequencing Centre, University of Copenhagen. To compensate for the low complexity in 5' ends of the CAGE libraries, 30% Phi-X spike-ins were added to each sequencing lane, as recommended by Illumina. CAGE reads were assigned to their respective originating sample according to identically matching barcodes. Using the FASTX Toolkit (v0.0.13), assigned reads were: i) 5’-end trimmed to remove linker sequences (10+1 bp to account for the CAGE protocol G-bias at the first 5’ base), ii) 3’-end trimmed to a length of 25 bp and iii) filtered for a minimum sequencing quality (Phred score) of 30 in 50% of the bases. Trimmed reads were mapped using Bowtie (33) (version 0.12.7) with parameters –t –best –strata –v –k 10 -y –p 6 –phred33-quals –chunksmbs 512 –e 120 –q – un to ASM294v2.26. To obtain bp resolution CAGE Transcription Start Sites (CTSSs) (21), the number CAGE tag 5’-ends were counted for each genomic position. CTSSs coordinates were offset by 1 bp to account for the G-bias trimming. Only chromosomes I, II and III were used for analysis. Mapping statistics are available in **Table S1**.

### Quantification and annotation of TSSs and genes

For each library, CTSS counts were normalized into tag-per-million (TPM) values and the sum of TPM-values across all libraries were calculated for each base pair. Nearby CTSSs within 20 bp of each other were merged into Tag Clusters (TCs). Only CTSSs passing a TPM-threshold (0.04516233) were considered; this threshold was chosen to maximize the total number of TCs across the genome (as implemented in the CAGEfightR R-package version 0.3, https://bioconductor.org/packages/CAGEfightR/). Expression was quantified as the number of tags in each TC for each sample. TCs having > 1 TPM in at least 3 libraries were retained.

Transcript- and gene-model annotations (ASM294v2.26) were downloaded from PomBase as a GTF-file and imported as a TxDb object (34). Regions were extracted using the *transcriptsBy-* family of functions and genes were extracted using the genes-function. TCs were annotated based on their overlap with annotated transcripts using the hierarchical models shown in Fig. 2A: Promoters were defined as the +/-100 bp region around a PomBase TSS, while Proximal was defined as −1000 to −100 from annotated TSS. 5’ UTRs, 3’ UTRs and CDS regions were defined as in PomBase. Exonic refers to non-protein-coding exons, including lncRNAs. Antisense was defined as intragenic but on the opposite strand. To assess gene-level expression, TCs were aggregated by summing all counts within or −1000 upstream of annotated genes. In case a TC overlapped more than one gene, the gene with the nearest annotated TSS was chosen.

**Fig 2.**
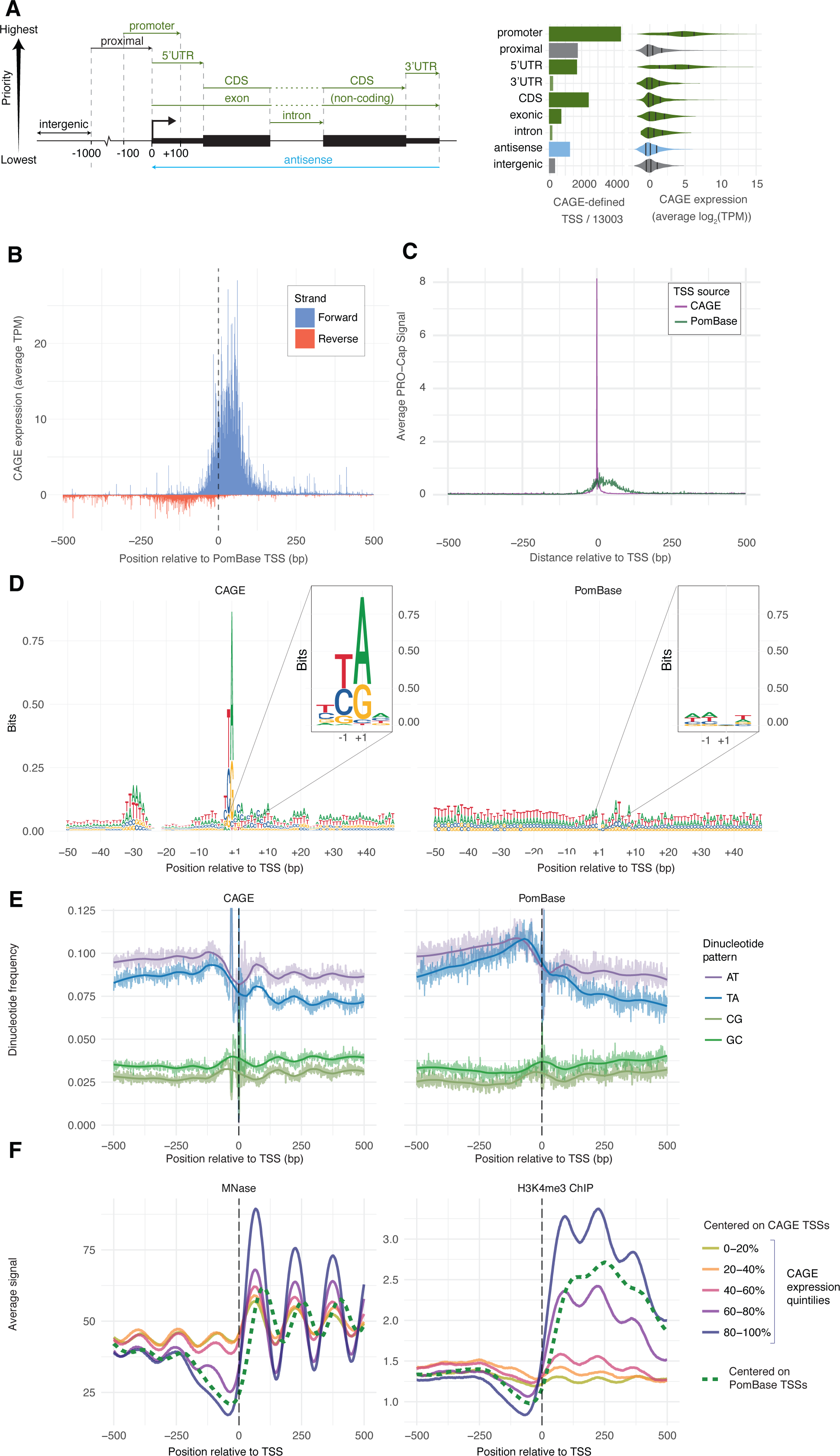
Properties of CAGE-defined TSS and comparison to PomBase annotation. **A: Location of CAGE-defined TSS vs. gene annotation.**Left panel: CAGE-defined TSSs were grouped into one of nine categories based on overlap with PomBase reference annotation (see schematic on top: categories are ordered by priority: in case a CAGE-defined TSS overlapped two or more categories, it was assigned to the one with the highest priority). Colors indicate whether regions overlap genes on same strand, antisense strand or are intergenic. Middle panel shows a bar plot counting the number of CAGE-defined TSS assigned to each category (from a total of 13003 expressed TSSs). Right panel shows the distribution of CAGE expression within each category (average TPM across all replicates and conditions) as violin plots, colored as above. Marks within distributions indicate 1^st^ quantile, median and 3^rd^ quantile.
**B: Comparison of CAGE and PomBase TSS locations.** X-axis show distance relative to annotated PomBase TSSs in bp. Y-axis shows average CAGE signal as TPM across all libraries: CAGE tags on the reverse strand in relation to the annotated TSS are shown as negative values (red), CAGE tags on forward strand are show as positive values (blue). Dotted line indicates PomBase TSS locations.
**C: Comparison of PRO-Cap signal at PomBase or CAGE-defined TSSs.** X-axis shows distance relative to TSSs defined by CAGE or PomBase. Y-axis shows the average PRO-Cap signal around CAGE-defined TSSs (purple) and annotated PomBase TSSs (green).
**D: Comparison of CAGE and PomBase promoter patterns.** Each panel shows a sequence logo of the +/-50 bp genomic region around TSS peaks defined by CAGE (left panel) or PomBase annotation (right panel). Y-axis shows information content in bits. Colors indicate DNA-bases. Insets show a zoom-in of the +/-2 region in each panel.
**E: Comparison of CAGE and PomBase promoter di-nucleotide frequencies.** Y-axis shows the frequency of di-nucleotides (as indicated by color) around TSSs defined by CAGE or PomBase; thick lines show loess regression trend lines. X-axes show distance relative to CAGE-defined TSSs (right panel) or PomBase-defined TSSs (right panel) in bp. Frequencies > 0-125 are cut.
**F: Comparison of MNAse-Seq and H3K4me3 ChIP-Seq signals anchored at CAGE or PomBase TSS.** X-axis indicates the distance from TSSs defined by CAGE or PomBase. Y-axis shows the average MNase-Seq or H3K4me3 ChIP-Seq signal, where larger values indicate highly positioned nucleosomes (left) or modified histones (right). Colors indicate whether signals are anchored on CAGE-defined TSSs (solid lines, stratified by expression quintiles) or PomBase-defined TSSs (dashed).

To make TSSs obtained from PomBase comparable to CAGE-defined TSSs in terms of expression, the same approach was used: CTSSs were quantified around PomBase TSSs (first bp of the 5’-UTR) +/-100 bp and were then TPM-filtered as above. CTSSs, TCs and transcript models were visualized using CAGEfightR and the Gviz R-package (35). TCs and bp-resolution tracks for each condition are deposited in the GEO database (GSE110976).

### Biological sequence analysis

The *getSeq-*function from the BSgenome package was used to extract sequences from the reference genome. Di- and trinucleotide frequencies were counted using *vmatchPattern-*function and sequence logos were made using the ggseqlogo package (36). TATA-Box and INR-motifs were obtained from JASPAR (37) via the JASPAR2016 R-package (38). The *motifScanScores-* function from the seqPattern R-package (http://bioconductor.org/packages/seqPattern/) was used to scan for motif occurrences. CpG frequency (CpG dinucleotides per bp) was calculated in a - 100/+100 window around TSS peaks.

### Average CAGE, MNase-seq, H3K4me3 ChIP-seq, PRO-Cap, PRO-Seq and NET-seq signal calculations

MNase-seq and H3K4me3 ChIP-Seq data was obtain from (39, 40) (accession numbers GSM1374060, GSE49575). Only wildtype strains were used in both cases. PRO-Cap and PRO-Seq signals were obtained from (41) (accession number GSE76142). NET-seq data was obtained from (42) (BAM-files were generously supplied by the authors). The GenomicAlignments package (34) was used to import and calculate the average signal.

In all cases, data was imported with rtracklayer and coverage calculated with the coverage-function from IRanges (34). CAGE, PRO-Cap, PRO-Seq, and NET-Seq profiles were plotted after removing the 1% most highly expressed sites, to remove the influence of a few extreme outliers.

For figure **3A-B,** only CAGE-defined TSSs at annotated promoters were used. For Fig. 3C and **Fig. S3C-D,** TSSs were selected based on the following criteria: i) the TSSs must have an upstream CAGE TSS within 250 bp on the opposite strand and ii) the TSSs must not be within 500 bp of an annotated TSS, on either strand. TSSs were grouped by the amount of NET-Seq sense signal in the nucleosome-depleted region (NDR), defined as −250:-50 bp relative to the CAGE-defined TSS peak.

**Figure 3.**
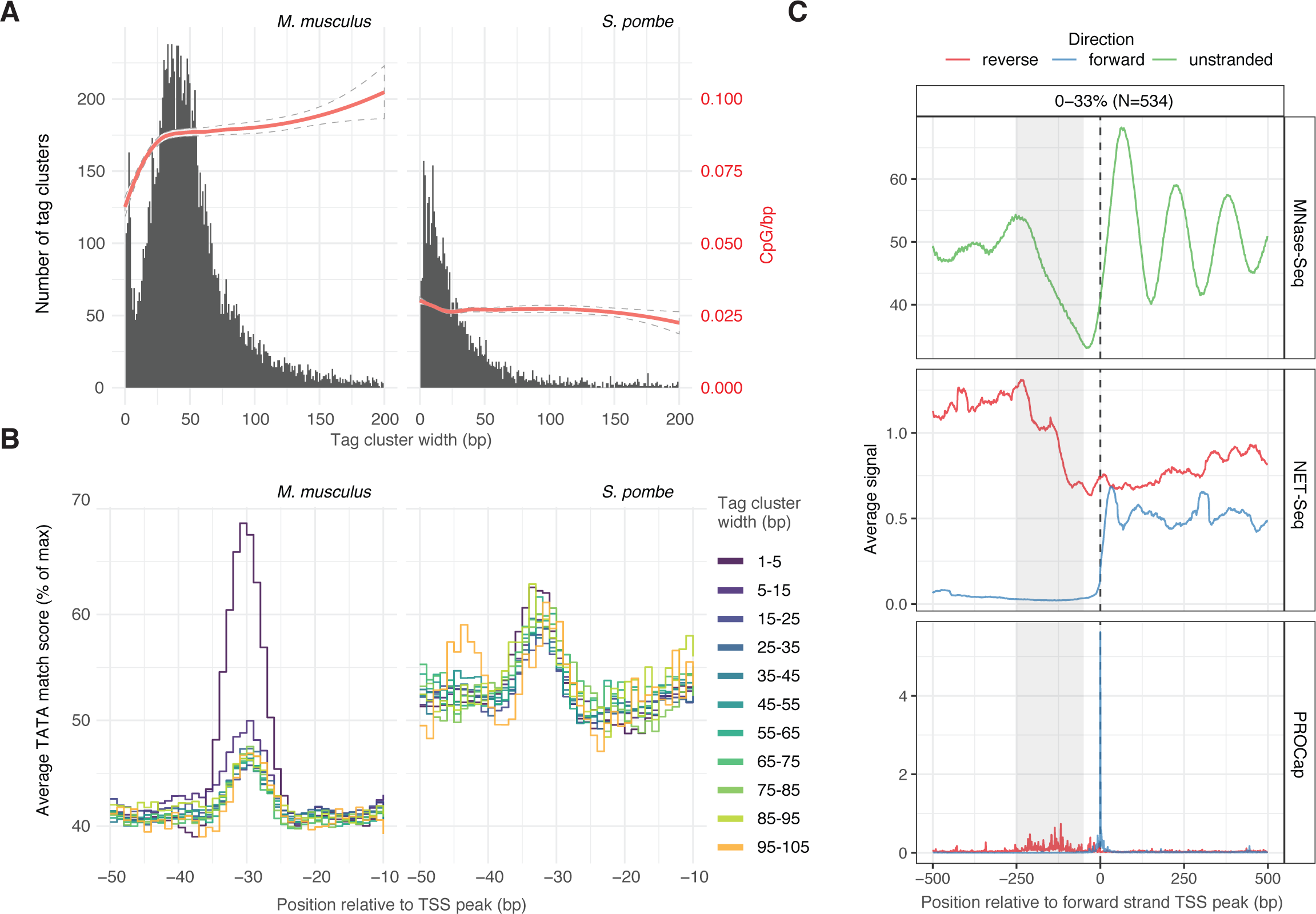
Analysis of TSS distribution shape and bidirectional transcription in S. pombe. **A: Relation between CAGE tag cluster width and CpG content.** X-axes show CAGE tag cluster width as defined in **Fig. S1A**. Left Y-axes and black histogram show the distribution of widths in five pooled *M. musculus* lung CAGE libraries (left panel) and this study (*S. pombe,* right panel). Red trend-lines and right Y-axes show the average number of CpG dinuclotides per bp (200 bp centered on TSS peaks). 95% confidence intervals are shown as dotted lines. Only CAGE-defined TSSs corresponding to annotated TSSs were analyzed.
**B: Relation between TATA-box occurrence and tag cluster width.** X-axes show the −50 to-10 region relative to TSS peaks. Y-axes show the average TATA-box motif match score (0-100% of maximal score). TSSs are stratified by their tag cluster width, indicated by line color. CAGE-defined TSSs from *M. musculus* and *S. pombe* are shown in left and right panel, respectively. Only CAGE-defined TSSs corresponding to annotated TSSs are analyzed.
**C: Analysis of bidirectional transcription initiation.**X-axes show distance relative to *S. pombe* CAGE-defined TSSs in bp. Each panel shows a different assay (MNase-Seq, NET-Seq or PRO-Cap). Y-axis shows average signal, in forward (blue), reverse (red) direction or unstranded (green) relative to the TSS. TSSs analyzed were selected to have bidirectional transcription based on CAGE, and subsequently filtered to only retain those with little sense NET-Seq transcription in the NDR-region (highlighted in grey). **Fig. S3C-D** shows TSSs with higher NET-Seq signal in the NDR region.

### Comparison with TSS atlases from Li *et al* and Eser *et al.*

Eser *et al* (43) TSSs were obtained from the transcriptional units from the supplementary material of the paper by selecting the first 5’-end position as the TSSs. Li *et al* (44) CTSSs and TCs were graciously provided by the authors. We retained Li *et al* TCs with >1 TPM and recalculated TC peaks similar to above to obtain TSSs.

### Comparison with mammalian TSSs

We obtained CTSSs from five control replicates for CAGE assays on mouse lung from (45). As these libraries were prepared in the same lab as the *S. pombe* libraries presented here and analyzed using the same pipeline, they are highly comparable. TC widths were calculated as the distance between the two positions in the TC corresponding to the 10^th^ and 90^th^ percentile of the pooled TPM across all libraries.

### Differential expression analysis and GO-term enrichment

All analysis was carried out using the edgeR (46) and limma (47) R-packages. The same analysis was run for TSSs and for genes. Initial normalization using the TMM-method from edgeR revealed a consistent shift in the overall distribution of expression values in the Nitro-EMM libraries. This could either reflect a large global shift in transcriptional output or an artifact of the additional RNA-purification steps in these samples (see above). Subsequent analysis using only TMM-normalization revealed a large number of up-regulated compared to down-regulated TSSs as well as a global shift in fold changes for highly expressed genes. For the final analysis we therefore chose the conservative approach of also normalizing expression using the cyclicloess-method prior to voom-transformation to model the mean variance relationship inherent to count data. Heat maps are based on the dual-normalized expression values. Multidimensional scaling (MDS) plots were made using the *plotMDS*-function from the limma package using pairwise Euclidian distances between the top 1000 genes/TSSs (top=1000) and otherwise standard settings.

We modeled expression using ~treatment+medium with Ctrl-EMM libraries as reference levels. We used robust estimation at the empirical Bayes stage and used the treat method to test against a minimum required log_2_ fold change of 0.5 (corresponding to ~41% higher or lower expression between states). Resulting *P*-values were corrected for multiple testing using Benjamini-Hochberg. **Fig. S4 and Fig. S5E-J** show diagnostic plots for the differential expression analysis on TSS and gene level.

Housekeeping candidates were identified using an F-test across all coefficients, with resulting *P*-values corrected for multiple testing using Benjamini-Hochberg. Biological Coefficient of Variations (BCVs) were calculated using the *estimateDisp-*function with robust=TRUE from the edgeR package. Genes with low BCV were identified by splitting TSSs/genes into 25 bins based on average expression, and selecting the 10% TSSs/genes with the lowest BCV.

Differential transcript usage was assessed using the diffSplice-approach, with the Simes-method to aggregate results at the gene level. Resulting *P*-values were corrected for multiple testing using the Benjamini-Hochberg method.

GO-terms were downloaded from the PomBase, and the kegga-function was used to test for enrichment, with mean expression used as the covariate. Resulting *P*-values were corrected for multiple testing using Benjamini-Hochberg.

### Comparison with previously defined CESR response and alternative 5’ UTR TSSs

CESR genes were downloaded from (4) (http://bahlerweb.cs.ucl.ac.uk/projects/stress/); 62/68 genes for the CESR Up/Down response could be matched to the CAGE data based on gene IDs. Log_2_ fold changes estimated with limma-voom were plotted for each CESR gene. Genes utilizing alternative 5’-UTR usage was obtained from (48). 24/28 genes could be matched by gene IDs. As the original data set did not provide coordinates for novel TSSs, we compared our gene-level differential transcript usage results from limma-voom with diffSplice to investigate agreements.

### Motif enrichment analysis in promoter regions

Promoter region DNA sequences for TSSs were extracted as sequences −300 to +100 of TSS peaks (**Fig. S1A**) via BSgenome and *getSeq* as described above. We used Homer (49) in *denovo* pattern discovery mode to detect enriched motifs in each differentially expressed set compared to a background set consisting of TSSs not part of any differentially expressed set (default setting except size=given, S=9, p=40 and mset=yeast). Because Homer works without prior knowledge of DNA-binding motifs, identified motifs may be novel patterns or correspond to binding preferences of known transcription factors. To link identified motifs with known motifs, we compared them to Homer’s database of S. *cerevisiae* motifs. The top 5 enriched *de-novo* motifs with a foreground set frequency of >0.9% were plotted using ggseqlogo. *S. cerevisiae* motifs was also used to perform known motif enrichment for the same sets (**Dataset S1**).

### Single Nucleotide Variation analysis

Genetic variation data was downloaded from EnsemblFungi (50). Only Single Nucleotide Variations (“TSA=SNV;Jeffares_SNPs”) were used for analysis. IRanges (34) were used to calculate the average number of SNVs per basepair in promoter regions (TSS peaks −300:+100 base pairs). GC-content and TATA-box positions were calculated as above. Background single nucleotide variation (SNV) frequencies were calculated as the total number of SNVs overlapping CDS regions divided by the total size in basepairs of all CDS regions. Filtering TSSs to only include TSSs at annotated promoters more than 500 bp distant from other such TSSs did not affect the results (data not shown).

The number of SNVs within promoter regions as a function of differential expression status and annotation category was modeled using negative binomial (NB) regression (due to overdispersion compared to a Poisson distribution) using the *glm.nb-*function from the MASS R-package (51). Exponentiated effect estimates and confidence intervals were extracted using the *tidy-function* from the broom R-package (https://CRAN.R-project.org/package=broom). Including average TSS expression in the model did not have any major effects (data not shown).

### Protein domains in genomic space

Pfam protein domains (52) were obtained from PomBase. Amino acid positions were multiplied by 3 and mapped back to genome using the *mapFromTranscripts-function* from GenomicFeatures (34). Domain disruptive TSSs were defined as TSSs that either: a) overlapped a domain or b) were downstream of a domain within the same gene.

### Northern Blot Analysis

Standard Northern analysis was performed on the RNA samples used for CAGE analysis. The hybridization probe was obtained by labelling a 270 bp PCR fragment from exon 5 of *cds1,* generated with the primers Cds1-F: ttactgcgtctattccttttg and Cds1-R: cgaagaattgagctgttcg.

### Western Blot analysis

Protein extracts from a Cds1-HA tagged strain and a wild type control treated as above (Ctrl – EMM, Nitro-EMM, Heat-YES and Ctrl-YES for both strains) were made by the TCA precipitation method. Extracts were fractionated by SDS-PAGE. After semi-dry transfer to a nitrocellulose membrane, Cds1 was detected using primary anti-HA (12CA5) monoclonal antibodies and HRP-coupled secondary anti-mouse antibodies.

## Results

### A TSS atlas for *S. pombe*

We prepared three biological replicates of *S. pombe* cultures growing under five different conditions: cells growing exponentially in EMM2 medium (designated as “Ctrl-EMM”), cells growing exponentially in YES (“Ctrl-YES”), heat-shocked cells growing in YES (“Heat-YES”), nitrogen starved cells growing in EMM2 (“Nitro-EMM”) and hydrogen peroxide-stressed cells growing in YES (“H_2_O_2_-YES”) (Fig. 1B). After RNA purification, we constructed three CAGE libraries per condition, corresponding to each biological replicate (the CAGE method is outlined in Fig. 1A). The average yield was 16.9 million uniquely mapping tags to the ASM294v2.26 assembly (**Table S1**).

Mapped CAGE tags closely located to each other on the same strand were grouped into Tag Clusters (TCs), similarly to (53). The expression of TCs across libraries were assessed as the number of CAGE tags within each cluster, normalized by sequencing depth in the respective library as Tags Per Million (TPM). We retained only TCs that had at least 1 TPM in three or more libraries. After cutoffs, we detected 13,003 TCs. For simplicity, we will refer to these TCs as “CAGE-defined TSSs”. For each of these 13,003 CAGE-defined TSSs, we identified the “TSS peak” as the single bp position with the highest TPM value across all libraries (**Fig. S1A)**.

In order to interpret the activity of CAGE-defined TSSs, it is useful to annotate them in reference to known transcripts and gene models. As an example, Fig. 1C shows the annotated TSSs of the *use1* and *pep7* genes, which are positioned bidirectionally ~1kb apart. Both annotated TSSs were detected by CAGE in all conditions, albeit consistently slightly downstream of the annotated TSSs (see below for a systematical investigation of this phenomenon). However, an unannotated TSS for *pep7* was detected 106 bp downstream of the PomBase-annotated TSSs. Unlike the annotated TSS, this TSS was only strongly expressed in H_2_O_2_-YES libraries. It is thus an H_2_O_2_-YES-specific novel alternative TSS. We show genome-browser examples of key stress response genes in **Fig. S1B-D**, and a more complex locus with multiple TSSs and transcripts in **Fig. S1E**.

### CAGE-defined TSSs refine PomBase TSS annotations

To systematically investigate the overlap with existing gene models, each CAGE-defined TSS was annotated into one of nine categories, depending on their overlap with transcript models as defined by PomBase (Fig. 2A, left). CAGE-defined TSS overlapped PomBase-annotated promoter regions (+/-100 bp from the start of PomBase annotated TSSs) in 34% of cases (Fig. 2A, right). Remaining CAGE-defined TSSs (8590 TSSs) commonly overlapped the proximal upstream region of genes (up to 1 kbp upstream of PomBase annotated TSSs, *14%*), 5’-UTRs *(13%),* and coding exons (CDS) (18%). Notably, CAGE-defined TSSs overlapping annotated promoters or 5’-UTRs were generally more highly expressed than CAGE-defined TSSs within other categories. This reflects observations in mouse and human (e.g. (45, 53)) and indicates that novel TSSs on average are less expressed, although clear exceptions to this rule exist (e.g. Fig. 1C).

We noted that while CAGE-defined TSSs often overlapped the regions around PomBase-annotated gene TSSs, the peak (the position having the most CAGE tags) of the corresponding CAGE tag cluster was often located slightly downstream of PomBase TSSs (exemplified in Fig. 1C). Plotting the average CAGE signal around PomBase-defined TSSs confirmed the existence of a small but consistent shift, where the majority of CAGE tags fell 50-75 bp downstream of annotated TSSs (Fig. 2B and **Fig. S2A**). This systematic difference could either be due to systematic overestimation of gene lengths in the PomBase annotation, or a systematic bias in our CAGE data. We reasoned that the most correct TSS set should better recapitulate known biology of gene regulation at DNA-sequence and chromatin levels.

First we compared our data to an independent PRO-Cap dataset (41). Like CAGE, PRO-Cap is based on sequencing of capped 5’-ends but captures nascent RNA. CAGE TSSs coincided with extremely high and focused PRO-Cap peaks (Fig. 2C) while most PRO-Cap signal was dispersed downstream of PomBase TSSs, consistent with the shift observed in the CAGE data (Fig. 2B). Additionally, NET-Seq (42) and PRO-Seq (41) data, capturing 3’-ends of nascent RNAs within RNA Polymerase II (RNAPII), showed peaks +30 (NET-Seq) and ~+55 (PRO-Seq) (**Fig. S2B**), likely corresponding to poised/stalled RNAPII that initiated transcription at the CAGE-defined TSSs.

Second, we investigated sequence content around TSSs. Previous experiments have shown that TATA and INR core promoter elements have a strong positional preference around TSS (~-32/-25 for TATA and immediately at the TSS for INR) when present (54, 55). A sequence logo (56) constructed by aligning the DNA sequence +/-50bp around all peaks of CAGE-defined TSS (Fig. 2D, left) showed enrichment of T/A-rich sequences at −32 to −25 and a strong pyrimidine-purine dinucleotide enrichment at −1/+1. These correspond to the TATA box and the central part of the INR element, and this pattern was highly similar to corresponding analysis in human and mouse (21, 55). While this pattern was present across CAGE-defined TSSs in all annotation categories as defined above **(Fig. S2C)**, it was not detected when constructing a sequence logo based on PomBase-defined TSSs (Fig. 2D, right). Dinucleotide analysis in the same regions showed a 150-bp cyclical pattern with higher AT/TA frequency downstream of CAGE TSSs (Fig. 2E, left). A similar, but weaker, pattern was present when centering on PomBase TSSs (Fig. 2E, right). Nucleosome structure in *S. pombe* is highly DNA sequence dependent, determined by relative frequencies of CG and TA dinucleotides (57–59). We reasoned that if the 150-bp cyclical AT-rich patterns reflected nucleosomal placement, then nucleosome alignment should be stronger and better when positioned by CAGE data compared to PomBase. To test this, we analyzed nucleosomal positions by MNase-Seq (micrococcal nuclease digestion followed by sequencing) in Ctrl-EMM cells from (39). Indeed, MNase-Seq signal displayed the same cyclic pattern downstream of CAGE-based TSSs as predicted from dinucleotide frequencies, where the amplitude, but not phase, correlated with the CAGE expression strength and the position of the +1 nucleosome edge was adjacent to the CAGE TSS (Fig. 2F, left and **Fig S2B**, top right). H3K4me3 ChIP-seq (40) signals from corresponding cells showed similar patterns (Fig. 2F, right, and **Fig S2B**, bottom right).

Finally we compared our CAGE data with two previous TSS atlases for *S. pombe*: Li *et al* (44) based on a single CAGE library and Eser *et al* (43) based on two RNA-Seq libraries. Compared to both these resources, our CAGE-based TSS atlas contained thousands of novel TSSs **(Fig.S2D)**. This likely reflected a) our much larger sample size and higher library depth and b) that our atlas covered a large range of environmental conditions. The majority of Li *et al* and Eser *et al* TSSs were confirmed by our CAGE-based TSS data **(Fig. S2E)**. Since the Li *et al* TSS atlas was also based on CAGE, there was high agreement of precise TSS locations with our set **(Fig. S2E, bottom)**, while Eser *et al* TSSs showed a systematic downstream shift in TSS positions **(Fig. S2E, top)**. This is not surprising given the random shearing and RNA length selection in the RNA-Seq protocol, which makes RNA-Seq prone to underestimating mRNA lengths. See **Table S2** for details on all TSSs and their relation to PomBase, Li *et al* and Eser *et al.*

Together, these observations (nascent RNAs and steady-state RNA approaches for locating TSSs, sequence content, nucleosome occupancy and histone modifications) strongly indicate that CAGE-defined TSSs are more accurate than current PomBase annotation, and greatly expands previous systematic attempts at locating TSSs. Thus, our data do not only increase the number of known TSSs, and thereby promoters of *S. pombe,* but additionally refines the positional accuracy of existing genome-wide data sets. The latter is important for any study relying on nucleotide-resolution TSS-anchored analysis and visualization; including profiling of histone marks, transcriptional activity, initiation and elongation. Our data thereby constitute a valuable new source of information for investigating *S. pombe* transcriptional regulation and chromatin biology.

### *S.pombe* promoters lack broad TSS distributions but a subset support bidirectional transcription initiation

One of the main advantages of *S. pombe* compared to *S. cerevisiae* as a model organism is its higher similarity to mammalian genomes in terms of gene regulation and chromatin biology. It was therefore important to assess whether core promoter features in mammalian genomes also were present in *S. pombe.* In particular, we compared the shape of TSS distributions, the prevalence of bidirectional transcription initiation and the relation between TSSs and nucleosomes in *S. pombe* vs. mammalian genomes.

In mammals, promoters can be distinguished by their distribution of TSS into “broad” and “sharp” classes. Sharply defined TSS distributions, where CAGE tags are focused on one or a few nearby nucleotides, are associated with TATA boxes and cell/tissue-specific transcription, while broader TSS distributions are often overlapping CpG islands and are associated with more ubiquitous expression (reviewed in (60)). Notably, in mammals, broad TSS distributions are more common than sharp. We measured the broadness of nearby TSSs as the width of CAGE tag clusters **(see Fig. S1A)** in *S. pombe* and a set of representative mammalian CAGE libraries produced and processed in the same way as the *S. pombe* libraries presented here (five pooled *M. musculus* lung CAGE libraries (45)).

While both *S. pombe* and *M. musculus* had CAGE tag cluster widths spanning from a single to hundreds of bps, *S. pombe* did not show the same bimodal distribution of tag cluster widths found in mouse; in particular, 30-50 bp wide tag clusters were common in *M. musculus* but not in S. *pombe.* This observation was correlated with the underlying DNA sequence content: while *M. musculus* tag cluster width, as previously reported, was positively correlated with CpG dinucleotide content (Fig. 3A), this relation was not present in *S. pombe,* perhaps related to the allover low CpG dinucleotide content in *S. pombe* promoters **Fig. S3A)**. Similarly, in *M. musculus* we observed a clear tendency for stronger TATA-box matches upstream of the peaks of sharp TSS distributions (width 1-5 bp); this pattern was absent in *S. pombe* (Fig. 3B); on the other hand, *S. pombe* tag clusters had, regardless of their width, average TATA match scores as high as sharp *M. musculus* tag clusters. This is likely partially due to the overall higher T/A content in *S. pombe* promoters. INR motif scores showed no relation to tag cluster width in either species (*Fig. S3B*). Overall, our results imply that *S. pombe* does not display broad TSS distributions as frequently as mammals, which may be due to the overall higher TA content and, in extension, higher TATA-box content.

Recent studies have shown that transcription in mammals initiates at both edges of the nucleosome depleted regions (NDR), producing mRNAs in the sense direction and typically short and exosome-sensitive RNAs in the reverse direction. In mammals, such reverse strand transcripts have been termed PROMPTS or uaTSS (61–63). Similar observations have been made in *S. cerevisiae,* where the transcripts are called CUTs or SUTs (reviewed in (61)), where CUTs, but not SUTs, are primary exosome targets. In *S. pombe,* the presence of PROMPTs has been debated: a study based on tiling-arrays detected PROMPT-like transcripts (62), while a recent study using nascent RNA sequencing (41) indicated that PROMPTs are uncommon. A third study showed that transcription of PROMPTs may occur but is repressed by the Spt5 protein (63).

To investigate this question in light of our CAGE TSS data, we re-analyzed recent nascent RNA sequencing data (NET-Seq, PRO-Cap and PRO-Seq) (41, 42). When anchoring upon all CAGE-defined TSSs, there was no overall pattern of bidirectional transcription in any of the three assays (data not shown), supporting the conclusion drawn in (41). However, through manual observation, we observed many cases where a CAGE-defined TSSs had a corresponding nearby bidirectional TSS (for example, see **Fig. S2E**). To see if such cases were supported by nascent transcription assays, we identified all CAGE-defined TSSs that also had upstream CAGE signal on the opposite strand within 250 bp. Because the *S. pombe* genome is gene-dense, we filtered cases having one or more additional annotated TSS within 500 bp on any strand.

Many if these CAGE TSSs showed reverse strand signal in PRO-cap, NET-seq and PRO-seq consistent with transcription initiation at the edges of the NDR, mirroring the properties of PROMPTs in mammals, but the signal was inversely correlated with the amount of sense transcription in the NDR, as measured by NET-Seq (Fig. 3C, **Fig. S3C**). A possible explanation for this observation is that read-through transcription from upstream genes on the same strand suppresses transcription initiation on the reverse strand, although the mechanism is unclear. Indeed, we generally observed stronger PROMPT transcription at lowly expressed forward strand CAGE-defined TSSs near or upstream of annotated promoters, while highly expressed forward strand TSSs in coding regions or 3’-UTR only showed weak PROMPT transcription (**Fig. S3D**).

In conclusion, the above analyses indicate that *S. pombe* TSSs share some, but not all mammalian properties: *S. pombe* promoters do not exhibit a clear distinction between broad and sharp TSS distributions, *S. pombe* TSSs are placed next to nucleosome boundaries but only a subset of them seem to support bidirectional initiation. Such cases typically corresponded to TSSs with limited read-through from other transcriptional units.

### Identification of stress-responding TSSs

In the above analysis, we treated all TSSs equally, even though they may be ubiquitously expressed across conditions, specifically expressed in one condition, or somewhere between these two extremes. Multidimensional scaling (MDS) analysis showed that libraries subjected to the same type of stress and medium clustered tightly together, indicating low biological variance within groups and consistent differences between groups (Fig. 4A). Nitro-EMM libraries, followed by Heat-YES, appeared the most different from Ctrl-EMM. Interestingly, we observed a clear difference between control libraries growing on the different growth media (Ctrl-EMM and Ctrl-YES), which was of comparable magnitude to the difference between Ctrl-EMM and H_2_O_2_-EMM. Thus, YES medium appears to alter the TSS landscape to an extent comparable to some stressors.

**Figure 4.**
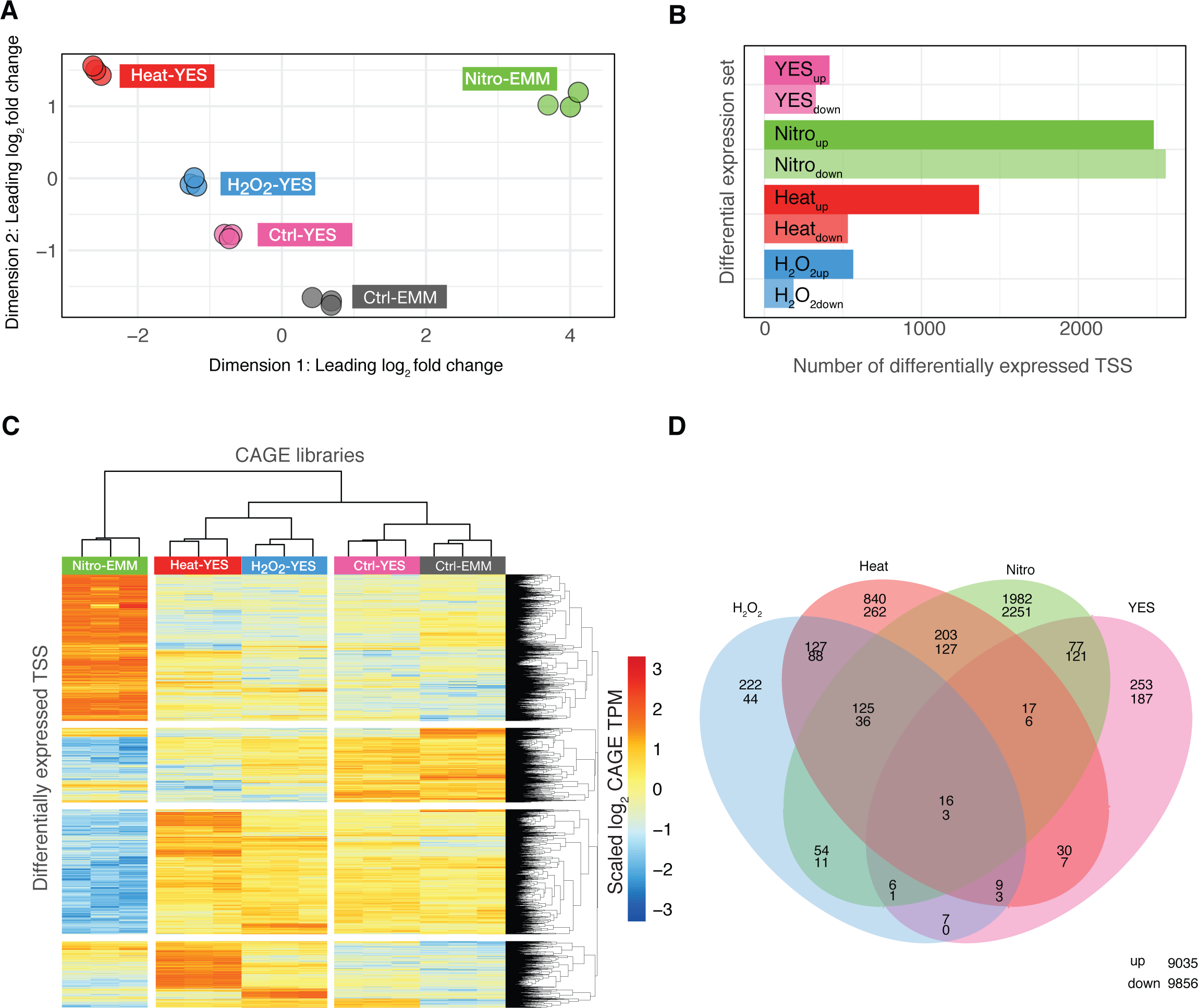
Differential expression of TSS in response to stress and media change. **A: Multi-dimensional scaling (MDS) plot of CAGE libraries.** MDS X-and Y-axes show the first two MDS dimensions. Each point corresponds to a CAGE library, colored by condition. Axes are scaled as leading log_2_ fold changes; the root-mean-squared average of the log_2_ fold changes of the top 1000 genes best separating each sample.
**B: Differential expression analysis on TSS level.** Bar plot shows the number of up-/down-regulated in each set of differentially expressed TSSs, corresponding to effects relative to the reference group (Ctrl-EMM) for the four main effects in the linear model. TSSs were considered significant if *FDR* < 0.05 and |log_2_ fold change| > 0.5, using limma-voom.
**C: Expression patterns of differentially expressed TSSs.** Heat map rows correspond to all differentially expressed TSSs in panel B. Columns correspond to all CAGE libraries, as indicated by color. Heat map shows the relative CAGE expression (log2(TPM) scaled to unit mean/variance across rows) for a given TSS and library.
**D: Overlap of differentially expressed TSS between treatments.** Venn diagram shows the overlap between sets of differentially expressed TSSs as defined in panel B. Number pairs within Venn diagram areas show the number of up- and down regulated TSSs (upper and lower number, respectively). Number in lower right corner are non-differentially expressed TSSs.

To identify individual TSSs that showed statistically significant changes in expression, we analyzed differential expression on TSS-level using the linear model framework implemented in the R-package limma (47) (*Fig. S4A-F*). The advantage of this approach, compared to multiple pairwise tests, is that the individual effects of media and different stressors can be estimated separately (e.g. the effect of YES vs EMM can be estimated jointly using all samples by controlling for the effects of stressors, rather than just comparing Ctrl-YES to Ctrl-EMM). Because of the high sequencing depth combined with the low variance within groups, we were able to detect even weak levels of differential expression across all ranges of TSS expression. However, even though subtle changes could reliably be detected, we reasoned that changes with a substantial fold change were the most biologically relevant. Thus, we only considered TSSs with an absolute log_2_ fold change ≥0.5 (corresponding to ~40% higher or lower expression between conditions) and FDR<0.05 to be significantly differentially expressed between two or more states. Using Ctrl-EMM as a baseline (or intercept in the linear model), we defined four sets of differentially expressed TSSs responding by either increasing or decreasing expression: Nitro_up_/Nitro_down_ for nitrogen starvation, Heat_up_/Heat_down_ for heat shock, H_2_O_2up_/ H_2_O_2down_ for hydrogen peroxide stress and YES_up_/YES_down_ for YES media (**Table S3**). The results recapitulated the MDS plot: the Nitro_up_ and Nitro_down_ sets had the highest number of differentially expressed TSSs, followed by Heat_up_/Heat_down_, YES_up_/YES_down_ and H_2_O_2up_/H_2_O_2down_ (Fig. 4B) Notably, while YES_up_/YES_down_ and Nitro_up_/Nitro_down_ had roughly as many up-as down-regulated TSSs, Heat_up_/Heat_down_ and H_2_O_2up_/ H_2_O_2down_ showed a substantially higher number of up-regulated vs. down-regulated TSSs.

The three stress conditions shared many differentially expressed TSSs, (Fig. 4C-D), indicating that a substantial part of stress response was generic. Notably, almost half (49%) of the 6375 detected *S. pombe* TSSs were differentially expressed in at least one stress condition (excluding YES response). The YES-response was generally composed of distinct TSSs compared to the stress response.

To investigate how the above results corresponded to known fission yeast biology, we related differential expression to existing Gene Ontology (GO) terms. Since GO-term annotations are defined at gene level, for this analysis, we measured gene expression by summing CAGE expression contribution from all CAGE-defined TSSs within a gene on the same strand as in (64), and used these expression values to perform gene-wise tests for differential gene expression in the same way as the TSS-level expression analysis above (**Fig. S5A-J**). Most differentially expressed gene sets were enriched for the expected GO terms (**Fig. S5H** shows the 10 top GO terms for each gene set, **Table S4** shows all terms), i.e. H_2_O_2up_ was enriched for “cellular response to oxidative stress”, Heat_up_ was enriched for “protein folding”, Nitro_up_ was enriched for “autophagy” and YES_down_ was enriched for “thiamine biosynthetic process”. We observed an enrichment of GO-terms related to meiosis in genes upregulated after nitrogen starvation (e.g. “conjugation with cellular fusion” and “meiosis 1”). This is expected, as sexual development in *S. pombe* is intimately linked to nitrogen starvation (65–67), and nitrogen starvation in EMM2 medium is particularly efficient in inducing this response. The combined activation of genes related to sexual development and stress response explains the overall larger number of differentially expressed TSSs/genes in Nitro_up_/Nitro_down_ compared to the other stressors.

The above results were in general agreement with a previous study of the *S. pombe* stress response based on microarrays (4), with a focus on defining a set of CESR genes. Despite our smaller range of stressors, we observed substantial overlap between the upregulated genes in our sets and the CESR-upregulated set: 38% (26/68) CESR genes were in H_2_O_2up_, Heat_up_, and Nitro_up_ gene sets, while only seven CESR genes were not present in any of our differentially up-regulated sets. Only 6/62 downregulated CESR genes were found in H_2_O_2down_, Heat_down_and Nitro_down_(**Fig. S6A**). This most likely reflects our stricter criteria for differential expression, since only five genes showed a different direction of differential expression in H_2_O_2up_/H_2_O_2down_, Heat_up_/Heat_down_compared to (4) (**Fig. S6B)**

Since we found that almost half of all TSSs were changing due to one or more stressors, we also wished to identify TSSs and genes that were not changing in expression across any stress conditions. To select such TSSs or genes we required an *FDR* >0.05 in any comparison between conditions (using an F-test against a log_2_ fold change different from 0), resulting in 19% (2530) of TSSs and 16% (950) of genes. These genes were enriched for GO-terms related to protein transport and the Golgi-apparatus, e.g. “intracellular protein transport” and “ER to Golgi vesicle-mediated transport” **(Fig. S7A**). TSSs/genes showing no differential expression between a wide range of conditions would be useful as candidates for “housekeeping TSSs/genes”, and thereby good references, e.g. used for qPCR controls. We reasoned that ideal candidates should show low variance of expression across and within conditions. Because variance is correlated with expression strength, we required candidates to show low variance of expression given their expression levels, which resulted in a list of 93 TSS and 49 housekeeping gene candidates (**Table S5**). We also looked specifically at a set of commonly used housekeeping genes: act1, adh1, atb2, cdc2, gad8, leu 1, ura4. While all these genes showed low variance of expression, some of them also showed differential expression between conditions. In particular, all housekeeping genes except act1 and cdc2 exhibited a high fold change in response to nitrogen starvation. Only cdc2 did not show differential expression in any comparison **(Fig. S7B-C)**. This finding implies that careful selection of reference genes/TSSs is important.

### Stress responding TSSs provide insights into *S. pombe* gene regulation

An important question how the various stress responses identified above are regulated. Because CAGE data enables both accurate localization of TSSs, and precise quantification of TSS usage across conditions, it is ideally suited for prediction of transcription factor binding sites (TFBSs) responsible for stress-specific gene regulation in upstream promoter regions.

We used HOMER (49) to identity enriched DNA motifs in the promoters around TSSs up- or down-regulated by each stressor (defined as the −300 to +100 bp around TSS peaks). Because there is no comprehensive resource of transcription factor motifs in fission yeast, we performed the analysis *de novo* (identifying over-represented motifs with no prior information). To determine whether any of the *de novo* motifs resembled known motifs, we compared them to HOMER’s database of motifs from *S. cerevisiae.*

As with GO terms, we observed motifs enriched in specific types of stress as well as motifs common across all stressors. In many cases, the enriched patterns reflected known *S. pombe* or *S. cerevisiae* biology (see Fig. 5 for a summary and full analysis in **Dataset.S1**). Heat_up_, H_2_O_2up_and Nitro_up_promoters all shared enrichment for the CST6 motif. *CST6* is a homolog of *Atf1* which is widely regarded as the central regulator of CESR (4, 68). Heat*up* promoters were highly enriched for a HSF1-like motif, while H_2_O_2up_promoters were enriched for a PAP1-like motif (*S. cerevisiae* homolog of *YAP1*). Heat shock factors and PAP1 are well-known transcription factors associated with the specific response to heat shock and hydrogen peroxide stress, respectively, in both *S. pombe* and *S. cerevisae* (73, 74).

**Figure 5.**
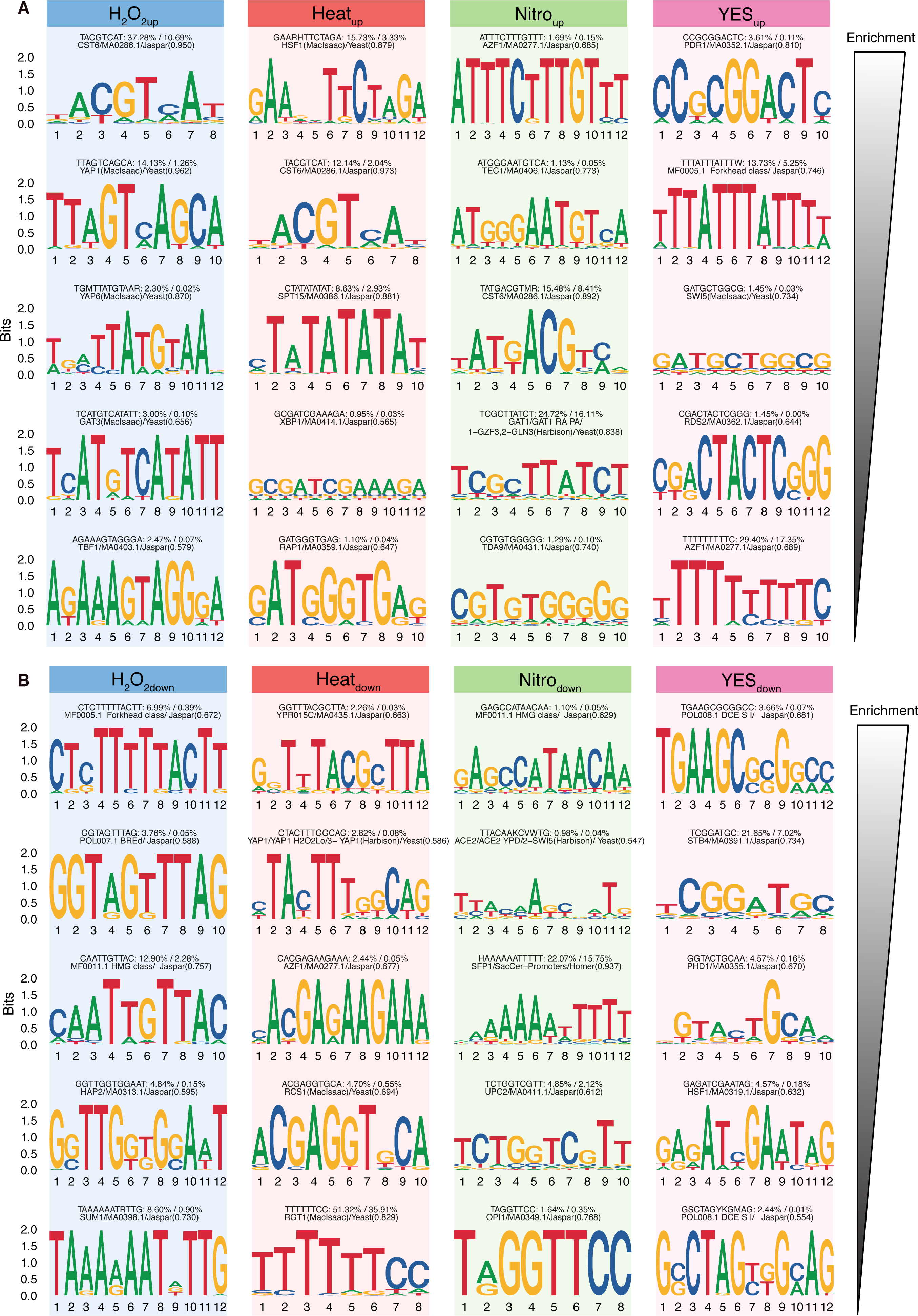
De-novo DNA motif enrichment in promoters of differentially expressed TSSs. **A: Enriched motifs in promoters of upregulated TSSs.** Columns show the top 5 enriched *denovo* motifs in the promoter regions of each set of differentially expressed TSSs (as in Fig. 4B), where the top motif is the most enriched. Motifs are shown as sequence logos, where Y-axis shows bit score and X-axis shows nucleotide positions in bp. Text above logos show consensus sequence, foreground / background frequency and closest matching known *S. cerevisae* motif (score in parenthesis indicated quality of match).
**B: Enriched motifs in promoters of downregulated TSSs.**Panels are arranged as in panel A, but show enriched motifs for promoters of downregulated TSSs.

Nitro_down_promoters were enriched for the SFP1 motif, while H_2_O_2down_was enriched for the similar SUM1 motif (while SFP1 was not found *de-novo* in heat_down_, a complimentary enrichment analysis using known *S. cerevisiae* motifs showed that SFP1 was enriched in heat_down_, H_2_O_2down_and Nitro_down_). SFP1 was previously implicated in regulating ribosomal gene expression (69), supporting the consistent enrichment of the “ribosome biogenesis” GO-term in corresponding down-regulated genes described above.

As discussed above, Nitro_up_were enriched for genes associated to sexual development in *S. pombe.* In agreement, the Ste11 motif (*AZF1* homolog) (66) was highly enriched in Nitro_up_as well as a GATA binding site (*GAT1* homolog). Ste11 and the GATA transcription factor Gaf1 have previously been shown to be regulated in a coordinated fashion in *S. pombe* (70).

YES_up_and YES_down_were enriched for different motifs when compared to the three types of stresses, reflecting an overall different regulation of this response. We are not aware of any comprehensive study of TFBSs involved in the response to YES-media, but our analysis suggests a possible role for several novel *S. pombe* TFBSs similar to known TFBSs in *S. cerevisae.* For example, YES_up_were enriched for a PDR1 motif and a general forkhead motif. PDR1 is annotated as specific to *S. cerevisiae,* while at least four forkhead-domain-containing genes exist in *S. pombe* (*fhl1, fkh2, mei4 and sep1*), which may have a role in cell adjustment to rich media. YES_down_promoters were enriched for GCR1, STB4 and PHD1 motifs. The GCR1/2 genes have not been shown to have *S. pombe* homologs, but regulate the glycolytic pathway in *S. cerevisiae* (71).

### Single nucleotide variation around *S. pombe* TSSs

TFBSs underlying the enriched motifs above were primarily detected in the −50 to −150 bp region relative to TSSs, suggesting that much of the regulatory signals for stress response may be located in this region. To investigate how this over-representation related to genetic variation across *S. pombe* populations, we investigated the overlap of CAGE-defined TSSs with available single nucleotide variations (SNVs) generated from comparison of multiple global*S. pombe* strains (50) (Fig. 6A). Overall, and consistent with (50), we observed a higher number of SNVs in intergenic regions (i.e. upstream of TSSs) compared to intragenic regions (downstream of TSSs). However, the precise location of the CAGE-defined TSSs revealed additional SNV patterns.

**Figure 6.**
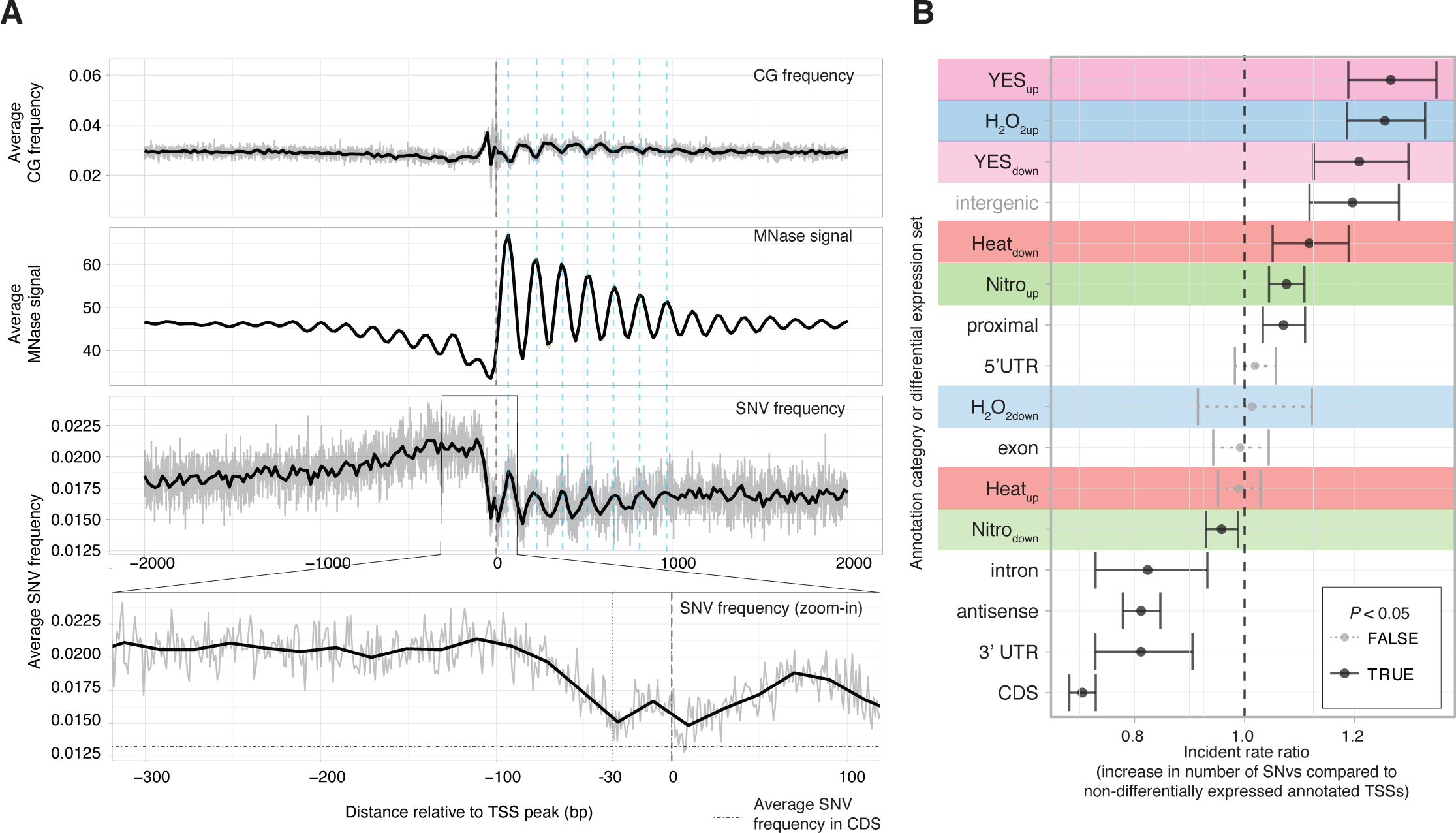
Single Nucleotide Variation (SNV) frequency around CAGE-defined TSSs. **A: SNVs around TSSs compared to CG-content and MNase-Seq signal.**X-axis in all panels shows distance from the CAGE-defined TSS peak in bp. Top panel shows average CG diucleotide frequency (grey lines show bp-level averages and black lines show a trend line, as in Fig. 2E). Middle panel shows average MNase-Seq signal, where larger values correspond to positioned nucleosomes (similar to Fig. 2F). The lower panel shows average number of SNVs, with zoom-in panel showing the −300 to +100 promoter region (grey lines show bp-level averages and black lines show a loess regression trend line). Vertical dashed teal lines indicate regions with high nucleosome occupancy. For the zoom-in panel, horizontal dot-dashed line indicates the average number of SNVs in coding exon regions as a reference, vertical dashed lines indicates the −30 position (typical TATA box position) and the 0 position (the TSS peak).
**B: Model of number of SNVs in promoters across annotation categories and differential expression sets.**Y-axis shows the different modeled effects, composed of different annotation categories (as in Fig. 2A) and the differential expression sets (as in Fig. 4B, also indicated by background colors). X-axis shows the incidence rate ratio: the estimated fold-change increase in the number of SNVs, compared to a non-differentially expressed TSS near a PomBase-annotated TSS. Points indicate estimates for the incident rate ratio for every modeled effect, whiskers indicate 95% confidence intervals. Whisker color indicates whether the *P*-value associated with an incident rate ratio is <0.05.

First, downstream of TSSs, SNV abundance was highly correlated to nucleosome occupancy (as defined by MNase-Seq data, see Fig. 2F) and anti-correlated to CG dinucleotide content (Fig. 6A). It is interesting to note that while CG dinucleotides in general are hyper-mutable, the TA-rich regions within the first 6-7 nucleosomes downstream of the TSS showed higher average SNVs than in CG-rich linker regions. A possible underlying mechanisms is that DNA with high nucleosome occupancy have higher mutation rates because nucleosomes block the access of the DNA repair machinery, as has been observed in human (72).

Second, SNV abundance was highest −400 to −100 bp upstream of TSSs, but rapidly declined towards two minima corresponding to the TATA box position at around −30 and the TSS at +1 (see zoom-in panel of Fig. 6A). The low frequency of observed SNVs around the TATA-box and TSS was similar to the SNVs frequency in coding exons. These results indicated that while the regions immediately surrounding the TSS and TATA-box were similar between *S. pombe* strains, the region upstream of the TSS was more variable: this was somewhat unexpected as this region overlapped most predicted TFBSs. To investigate whether this pattern was related to differential expression, we examined whether the regions around differentially expressed TSSs had more or less SNVs vs. non-differentially expressed TSSs. Specifically, we modeled the number of SNVs in the −300 to +100 region around the TSSs as a function of i) their differential expression (Heat*up*/Heat*down*, YES*up*/YES*down*, etc. as in Fig. 4B) and ii) their overlap with gene annotation (annotated TSS, 5’ UTR, coding sequence, etc., as in **xFig. 2A) using negative binomial regression (Fig. 6B). The advantage with this approach is that the effect of TSS location and differential expression status can be separated. Compared to a baseline consisting of non-differentially expressed TSSs near annotated promoters, intergenic TSSs (annotated as proximal or intergenic) had more SNVs, while intragenic TSSs (intron, antisense, 3’-UTR and CDS) had less (in agreement with Fig. 6A)**. Correcting for these gene location effects through the regression model, we found that some differentially expressed sets of TSSs were associated with substantial increases in the number of SNVs. In particular, YES_up_and YES_dowm_promoters had an increase in SNVs larger than the effect of being in an intergenic region. H_2_O_2up_, Heat_down_and Nitro_up_promoters were also enriched for SNVs, while H_2_O_2down_, Heat_up_and Nitro_down_were not enriched or depleted. We speculate that this increase in SNVs in differentially expressed promoters in particular could be related to positive selection or genetic drift of SNVs in these regions (see Discussion).

### Stress-specific activation of alternative TSSs

Alternative TSSs are common in multicellular organisms (29, 30, 73). Principally, genes may utilize alternative TSSs for at least three reasons: i) Having multiple TSSs or promoters may increase the overall expression of the gene in question; ii) If alternative TSSs are located within genes, their usage may produce RNAs that exclude coding exons resulting in truncated proteins, which may have a functional impact. iii) Multiple TSSs offer higher regulatory flexibility, where two TSS may respond to different stimuli or context due to different transcription factor binding sites around them.

While several studies have shown context-specific usage of alternative TSSs that may be disruptive and produce truncated proteins in *S. cerevisiae* (e.g. (74–77)), to our knowledge, no systematic attempts to profile alternative TSS usage in the *S. pombe* genome under different environmental conditions have been done. We reasoned that since CAGE can detect and quantify both known and novel TSS, our data set is well suited for detecting differential TSS usage within genes across stress conditions.

First, we investigated the occurrence of multiple TSS inside genes across all conditions. Around 54% of genes had >1 TSSs (Fig. 7A), indicating that multiple TSSs per gene is a widespread phenomenon in *S. pombe.* For many genes, a single dominant TSS might be expressed at much higher levels than other TSS(s) within the same gene. We therefore identified the dominant TSS within each gene (defined as the TSS with the highest overall expression across all libraries) and analyzed where dominant and non-dominant TSSs were located respective to annotation (Fig. 7B). For genes having >1 TSSs, the dominant TSS most often overlapped the annotated TSS region or 5’ UTRs, much like the location of TSSs with only a single TSS. Non-dominant TSSs were most commonly found in coding regions (CDS) or proximal upstream regions, a pattern consistent with the overall expression of TSSs in the same regions (Fig. 2A).

**Figure 7.**
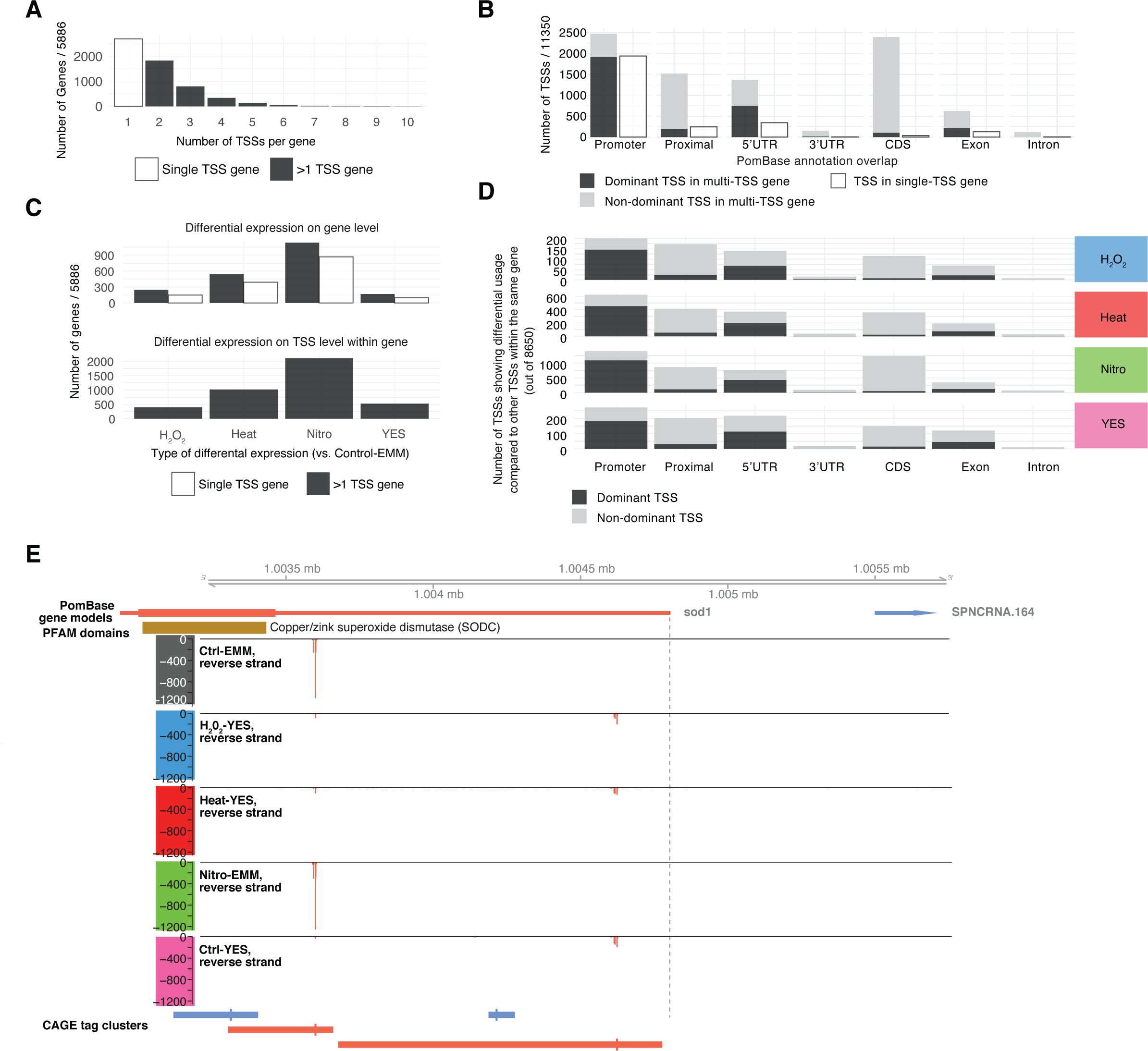
Differential TSS usage within genes. **A: Number of TSSs per gene.** X-axis indicates the number of TSSs per gene and Y-axis the number of genes. Colors indicate single-TSS genes (white) and multi-TSS genes (black).
**B: TSS structure compared to annotation and expression.** X-axis shows the annotation categories of intragenic TSSs (as in Fig. 2A), bar plots show the number of TSSs in each category, sub-divided into dominant (most highly expressed, black) and non-dominant TSSs within genes having >1 TSS (grey) and TSSs within single-TSS genes (white).
**C: Number of genes showing differential TSS usage compared to differential gene expression.**The top panel shows the total number of differentially expressed genes for each effect; genes are divided into those having one or more TSSs. The bottom panel shows the number of genes having at least one TSS with differential expression compared to the rest of TSSs within the gene. X-axes indicate effects and Y-axis indicates the number of genes.
**D: Differential TSS usage across different annotation categories.**Bar plots show the number of TSSs having differential expression compared to the rest of TSSs within the same gene, split by effect (rows, indicated by color), and PomBase annotation overlap (columns).
**E: Genome browser example of differential TSS usage in the sod1 locus.** Plot is organized as in Fig. 1C. See main text for details.

Next, we wanted to investigate how often multi-TSS genes used different TSS under different conditions. For each gene, we analyzed whether any of its TSSs showed a different response across conditions compared to the other TSSs in the same gene (using the diffSplice approach from limma, see Methods). Nitrogen starvation resulted in the highest number of genes showing differential TSS usage, followed by Heat, H_2_O_2_ and YES, mirroring the overall gene expression results (Fig. 7C-D). Overall, 77% percent of multi-TSS genes showed differential TSS expression compared to the gene’s other TSSs in at least one condition. It is thus common for genes to have at least two TSSs that are regulated differently in *S. pombe.* There was no clear preference for differential TSS usage in any specific part of genes (Fig. 7D) when comparing to the overall number TSSs in those categories (Fig. 7B), i.e. there was no enrichment (or depletion) of differential TSS usage in any particular part of genes compared to other TSSs. The largest number of differentially used TSSs was therefore observed in annotated promoter regions or 5’-UTRs. This pattern was consistent across all conditions.

Many genes showed both differential expression and differential TSS usage as defined above (**Fig. S8A**), indicating that these processes are coupled. We observed the same overall pattern in the top most expressed TSSs within genes (only TSSs making up at least 10 % of total gene expression in at least 3 libraries) (**Fig. S8B-D**).

In several cases, differential TSSs usage detected by CAGE verified previous single-gene studies. For example, we detected two TSSs in the 5’ UTR of the Superoxide Dismutase 1 (sod1) gene where most of the upstream TSS showed little change between conditions, but the downstream TSS (~1kb downstream of the annotated sod1 TSS) showed high expression in all EMM2 media conditions and low expression in YES-media conditions (Fig. 7E). Two *sod1* mRNA isoforms were previously identified (78): a shorter transcript that was gradually replaced by a larger transcript when cultures growing in YES entered stationary phase. The lengths of both transcripts detected in their Northern blot corresponded to the distance between the two TSSs detected by CAGE, suggesting that a switch from EMM2 to YES can create a similar change. The gpa1 locus is another example of differential expression of alternative TSSs **(Fig. S8E)**

We were also interested in the specific cases where differential TSS usage could lead to an altered protein product. We reasoned the most dramatic effect on the final protein product of a gene would be the exclusion or truncation of protein domains by the use of alternative TSSs. We identified TSSs that were located within or downstream protein domains defined in PomBase. Usage of these “disruptive” TSSs would give rise to either domain truncation or exclusion. We found that 64% of CDS TSSs, 56% of intronic TSSs and 75% of 3’-UTR TSSs were classified as disruptive. The average expression level of disruptive TSSs was low, and disruptive TSSs generally made low contributions to total gene expression (Fig. 8A). We detected 52 dominant disruptive TSSs, and 478 disruptive TSSs which contributed more than 10% of the expression of their host gene in at least three libraries. Of these TSSs, 356 were differentially used in at least one condition (Fig. 8B). This indicates that while alternative TSS usage affecting protein domain inclusion is relatively rare, there are clear cases of this phenomenon.

**Figure 8.**
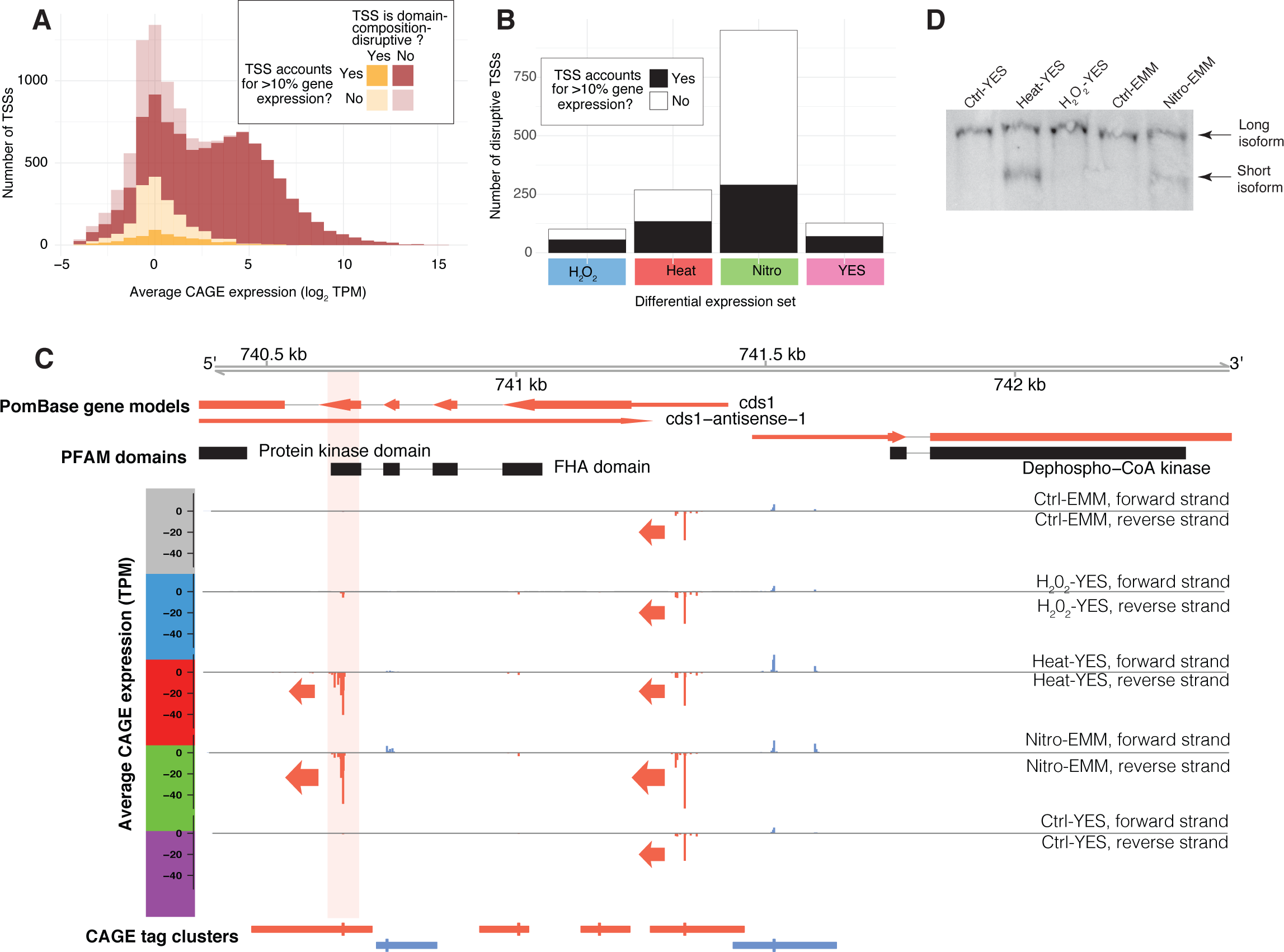
Candidates for alternative TSSs affecting domain composition. **A: Expression levels of domain composition-disruptive TSSs (TSSs in or downstream of protein domains):**Stacked histogram of TSS expression; X axis show log_2_(TPM), Y axis show number of TSSs. TSSs are split depending on whether the TSS disrupts domain structure and whether it contributes more than 10% of total gene expression in at least 3 libraries, as indicated by legend.
**B: Number of differentially used alternative TSSs potentially affecting domain composition:**X-axis shows the different conditions. Y-axis shows the number of differentially used TSSs with potential to disrupt domain composition. Color indicates whether the TSS contributes more than 10% of total gene expression in at least 3 libraries (See also **Fig. S8B-D**).
**C: Genome browser example of differential TSS usage in the*cds1* locus.**Plot is organized as in Fig. 1C, but also shows PFAM domains. TSSs downstream of, or overlapping, PFAM domains are highlighted with light red background. See main text for details.
**D: Northern analysis of*cdsi1* using a probe in exon 5.** The same RNA samples used for constructing the CAGE libraries were analysed by Northern blotting. The blot was hybridized to a probe in exon 5 of *cds1.* In addition to the full-length transcript present in all samples, the shorter *cds1* isoform, originating from the downstream TSS identified in panel C, is only evident in the Heat-YES and Nitro-EMM samples (lower bands).

The *cds1* gene represents an example of a domain-disruptive TSS (Fig. 8C). *cds1* had two highly expressed TSSs: one in the 5’ UTR which was active across all conditions, and one in the fourth protein coding exon, expressed only in Nitro-EMM and Heat-YES. The latter TSS would produce an RNA that cannot encode the forkhead domain encoded in the upstream exons. A previous study showed that this domain is important for the activation of Cds1 through binding to Mrc1 (79). We validated the Nitro-EMM and Heat-YES-specific expression of the shorter isoform using Northern blotting (Fig. 8D).

An importance caveat is that it is not possible based on CAGE data alone to determine if the transcript produced by an alternative TSS is translated into a stable protein product. While CAGE can be used as a hypothesis generator, protein-based experiments are needed to verify candidates for protein-altering alternative TSSs. Illustrating this caveat, a western blot for Cds1 after heat shock and nitrogen starvation did not detect a protein product corresponding to the Heat-YES/Nitro-EMM specific downstream TSS discussed above, despite the high RNA levels detected by CAGE (**Fig. S9A**). fus1 and arg11 genes are additional examples of domain-disruptive TSS **(Fig. S9B-C**).

## Discussion

Here we have used CAGE to define a comprehensive atlas of TSSs in *S. pombe* and their activity across a range of different growth conditions and stresses. We then compared our TSSs atlas to existing *S. pombe* annotation from PomBase, which is primarily based on RNA-Seq. We show that our TSS atlas can refine the location of known TSSs, including TSSs for protein coding genes, as well as more lowly expressed non-coding transcripts. This is in line with previous work (48), showing pervasive transcription of many unstable transcripts, including many anti-sense transcripts as exemplified in **Fig. S1E**.

Accurate TSSs are essential for studying chromatin biology, for which *S. pombe* is a widely used model organism. CAGE-defined TSSs had expected core promoter motifs at expected spacing, and showed a much better agreement with MNase-Seq and ChiP-Seq data for relating transcription initiation with nucleosome positioning and histone modifications respectively. We found that the typical peak distribution of TSSs in *S. pombe* promoters was considerably sharper than that of mammals, with a higher incidence of TATA boxes, perhaps due to the higher TA content in the *S. pombe* genome. As in vertebrates, TSSs are located next to nucleosome boundaries; on the other hand, only a subset of promoters seemed to support initiation of bidirectional transcription. This may be an effect of the closely located genes in *S. pombe,* as the strongest bidirectional TSSs were observed in cases with little read-through from other genes.

The only genome-wide TSS experiment previously reported for *S. pombe* is, to our knowledge, the single replicate CAGE experiment presented in (44). This study showed that TATA and INRs motifs in *S. pombe* are more similar to their vertebrate counterparts than *S. cerevisiae,* which our data also confirms. Li *et al* indicate that CAGE-based TSSs had much lower frequencies of TATA/INR patterns if they were not proximal to PomBase TSS; our data shows that even novel TSSs have TATA/INR patterns. This is likely an effect of our higher number of mapped CAGE tags and lower rRNA contamination vs Li *et al* (**Table S1**). Lastly Li *et al* found a novel motif upstream of a small subset of very sharply defined TSS distributions. Since our motif analysis was based on comparisons between environmental states rather than genomic background, it is not surprising we did not find the same pattern.

Our differential expression analysis on gene- and TSS-level shows that nearly half of TSSs are affected by stress response, with expression consistent with previous microarray studies (4). The overlap is noteworthy, since our study only measures RNA levels at a single time point following stress application, and thus cannot detect genes showing complex responses over time. Interestingly, we also observed a large transcriptional effect of YES-compared to EMM2-media, comparable in overall magnitude to some stresses. A similar phenomenon was reported in (48). The implication of this is that the two most commonly used growth media for *S. pombe* are not interchangeable, and thus, biological conclusions based on experiments performed on YES growing cells cannot necessarily be extrapolated to EMM2 conditions and vice versa.

This is also the first dedicated genome-wide study of differentially expressed alternative TSSs in *S. pombe* across stress conditions. We show that differential TSS usage is widespread in the S. *pombe* genome: 54% of genes utilized more than one TSS, and 77% percent of these showed differential alternative TSS usage in at least one condition. The large number of alternative TSSs reported here disagrees with a previous effort to detect genes with alternative TSS in 5’-UTRs using RNA-Seq (48), which identified only 30 genes. Using our data, we could confirm 23 of these cases for 28 genes found in both studies. The discrepancy in numbers is likely due to technical issues: since RNA-Seq is based on random fragmentation, it is challenging to identify distinct 5’-ends of full-length RNAs, as opposed to a dedicated technique like CAGE that specifically selects for them through cap-trapping. It is thus likely that *S. pombe* uses a large repertoire of alternative TSSs. Because multi-exonic genes, required for the generation of alternative splice isoforms, are uncommon in *S. pombe* (only 19.5% percent of genes have >2 exons), our data suggests that alternative TSSs are a greater source of transcript diversity than alternative splicing in *S. pombe,* unlike in mammals where the two processes seem be used to a similar extent (29, 80). We show that most differential TSS usage, where two or more TSSs for the same gene show different transcriptional patterns, involves TSSs located upstream of the annotated TSS or within the 5’-UTR. Differential regulation of such TSSs likely does not change the protein product of the resulting transcript but may confer regulatory flexibility because the two regions can evolve independently.

Binding of transcription factors to sites proximal to TSSs is a key process in gene regulation, and differential expression can be related to the accurate promoter regions by our TSS atlas. Using such regions, we identified likely TFBSs involved in the stress response, ranging from well-known stress-associated transcription factors to possible candidate transcription factors that may be preferentially used in YES or EMM2 media. We reasoned that the regulatory importance of these promoter regions may be related to their evolution. Indeed, we observed a strong conservation of the immediate TSS and TATA-box regions (comparable to that of coding sequence), while regions upstream of the TATA-box were more genetically variable between *S. pombe* populations. A simple explanation for this observed pattern is a lack of negative selection in the predominantly intergenic upstream regions, leading to a higher rate of genetic drift between *S. pombe* strains. However, we also observe that differentially expressed TSSs in many cases show more genetic variability, in particular YES-responding TSSs. Therefore, an alternative model is that while the relatively constrained CDS and core promoter regions are under negative selection, positive selection may be acting on genetic variants affecting transcription factor binding upstream of the TATA-box, thus giving rise to variation in gene regulation between *S. pombe* strains, rather than variation in the structure of genes themselves. Because YES-media is richer than EMM2, we speculate this higher diversity may be related to gradual adaptation to different nutritional environments for the different *S. pombe* strains, while the more acute stress (e.g. heat shock) response patterns may have a higher conservation of their regulatory elements.

While most of the work in this paper has focused on genome-level interpretation of the TSS atlas, we believe one of the main uses will be as a resource for detailed investigation of individual genes and processes. To make the TSS atlas as easily available as possible to researchers, it is available as a collection of genome browser tracks via the PomBase genome browser.

## Acknowledgements

We thank Carsten Skou Nielsen and Peter Brodersen for help with the Northern analysis. Work in the AS laboratory was supported by grants from the Lundbeck Foundation and the Novo Nordisk Foundation. Work in ON’s laboratory was funded by Villum Fonden. Work in the KE laboratory was supported by the Swedish Research Council and Cancerfonden.

## Author contributions

MT, AT, AA, YC, AS analyzed data. JL, MB made CAGE libraries. ON made *S. pombe* cultures and Northern blotting. CH performed Western blotting. MT, KE, AN, CW, AS interpreted results. MT, ON, AS wrote the paper with input from all authors.

## Data accession

CAGE raw data, as well as processed tag cluster and CTSS datasets have been deposited in the GEO database (accession number: GSE110976). The datasets, in summarized form, are also accessible through the PomBase genome browser (Under “Transcription Start Sites”) Supplementary figures and data sets are available at Figshare:

Supplementary Figures: https://doi.org/10.6084/m9.figshare.5977900
Supplementary tables: https://doi.org/10.6084/m9.figshare.5977846
Supplementary data set 1: https://doi.org/10.6084/m9.figshare.5977879

## Datasets

Dataset.1: Compressed archive of complete results of*de-novo* and known motif enrichment analysis with Homer. Each differentially expressed set has its own folder, containing a web page named “homerResults.html” for*de-novo* motif analysis and “knownResults.html” for known motif analysis, as well as position weight matrices for all motifs. For more information on the format see http://homer.ucsd.edu/homer/ngs/peakMotifs.html.

